# Tau alters global gene expression affecting chromatin in the early stages of Alzheimer’s disease

**DOI:** 10.1101/2023.11.17.567548

**Authors:** G. Siano, M. Varisco, M. Terrigno, C. Wang, Marco Groth, M.C. Galas, Jeroen J.M. Hoozemans, A. Cellerino, A. Cattaneo, C. Di Primio

## Abstract

Recent research revealed that Tau plays critical roles in various neuronal functions. Our previous work demonstrated that destabilization and nuclear delocalization of Tau can alter the expression of glutamatergic genes, thereby mediating early neuronal damage.

Upon analysing gene coexpression of AD temporal regions at different stages and the transcriptional output in differentiated neuroblastoma cells, we discovered that changes in Tau availability are linked to global alterations in gene expression that affect multiple neuronal pathways. Comparison with the human AD temporal region showed that the Tau-dependent modulation of gene expression closely resembles an intermediate stage of AD that precedes the definitive AD condition. Furthermore, we identified the chromatin remodelling pathway as being significantly affected by Tau in both our cellular model and AD brains, with reductions in heterochromatin markers.

We propose that Tau is able to globally affect the neuronal transcriptome and that its availability is linked to changes in gene expression during intermediate stages of AD development. In addition, we found that the epigenetic pathway is strongly affected by Tau during the progression of AD.

## Introduction

The microtubule (MT)-associated protein Tau is a key factor in the maintenance of neuronal homeostasis and cytoskeletal stability (Guo et al., 2017). During the early stages of tauopathies, Tau exhibits specific characteristics: its expression levels increase, its soluble fraction expands, and it progressively forms oligomers and aggregates that result in neuronal damage and cell death (Han et al., 2017; Kurbatskaya et al., 2016; Wang and Mandelkow, 2016).

Aggregates are commonly considered the main cause of neurodegeneration; however, the recently discovered Tau nuclear functions indicate a wider and more complex involvement in neuronal homeostasis changes in the very early stages of the disease (Siano et al., 2021; Sotiropoulos et al., 2017).

Tau binds chromatin and plays a relevant role in genomic stability, protecting DNA from damage and nucleolar organization under stress conditions (Colnaghi et al., 2020; Maina et al., 2018; Mansuroglu et al., 2016; Rossi et al., 2008; Sjöberg et al., 2006; Violet et al., 2014). Tau is involved in transposon activation, an event strongly correlated with Alzheimer’s disease (AD) (Guo et al., 2018; Sun et al., 2018). Tau also interacts with TRIM28, a nuclear factor that regulates chromatin remodelling and is associated with genomic instability in AD (Rousseaux et al., 2016). These findings suggest that the accumulation of Tau in the nuclear compartment might affect genomic organization and lead to neuronal damage.

We recently reported that nuclear Tau alters the expression of glutamatergic genes, thus causing toxic hyperexcitability in the early stages of AD. In contrast, aggregation prevents this mechanism, reducing glutamatergic deregulation (G. Siano et al., 2020; Siano et al., 2019b). These data have been further confirmed by a transcriptomic experiment in HEK cells in which *Montalbano et al.* identified approximately one hundred Tau-dependent altered genes involved in chromatin organization and remodelling, nuclear speckle structure and different pathology-related mechanisms, such as cytoskeletal organization and ubiquitin-related processes (Montalbano et al., 2021). The links among Tau, chromatin structure and stability and the role of Tau in modulating gene expression strongly suggest that these two processes might be connected.

Here, using a high-resolution transcriptomic approach, we investigated the Tau-dependent modulation of gene expression in a widely used human neuronal cellular model, the neuroblastoma SH-SY5Y line. We found that Tau induces global transcriptomic changes associated with early stages of AD and that one of the main altered pathways is chromatin remodelling, specifically histone modifications and heterochromatin organization. Our findings suggest that Tau alters the expression of several genes in the early stages and that the resulting modification of chromatin structure is a downstream event that may exacerbate the pathology.

## Results

### Longitudinal weighted gene coexpression network analysis (WGCNA) of AD brains reveals progressive alteration of transcriptomic pathways

To investigate the pathological mechanisms progressively altered during the progression of Alzheimer’s disease, we performed WGCNA on temporal lobe samples from healthy and AD human brains. WGCNA uses a distance matrix based on gene coregulation to identify networks of coexpressed genes called modules, which can later be mapped to Gene Ontology (GO) categories to infer their biological functions. Modules are built on all available data with an unbiased approach, and GO annotation is performed *a posteriori* on the whole module. In this way, the analysis is not limited to genes with known GO annotation: genes not included in the GO database can still be assigned to the module based on coregulation, and their function may be inferred from the GO of the module.

The coregulation networks also enable groups of concerted genes that include more than one GO category to be addressed. In addition, the changes in intramodular connectivity, both in the number of genes included in a module (i.e., the nodes) and in the strength of their connections (i.e., the edges), are highly informative for predicting the regulation point at which a pathway is altered.

For these analyses, we employed a published dataset, GSE84422 (GPL97 platform), containing data for AD brain samples structured into 3 groups: possible AD (posAD), probable AD (proAD) and definitive AD (defAD), representing early, intermediate and late phases, respectively (Wang et al., 2016). Healthy samples were analysed to build the reference, and 25 consensus modules were identified in control subjects. To follow the evolution of coregulated genes during the progression of the disease, the genes in proAD and defAD were assigned to these consensus modules (Fig 1A). Then, the conservation of reference modules in the proAD and defAD phases was measured via assessment of module preservation. To identify subsets of genes with the same trend, genes were clustered by their expression values at each stage with the K-means algorithm and Euclidean distance to preserve information on both the sign and magnitude of the log2 (fold change). By network analysis, we observed 2 modules that are not conserved in AD brains compared to healthy brains: the Pink and the Darkgrey modules (Fig 1B). In GO analysis of these modules, DNA binding and transcription regulation (Fig 1C) were featured among the functions listed in Table EV1 and Table EV2, suggesting that pathways associated with chromatin and gene expression might be altered during the progression of the disease.

**Figure 1.**
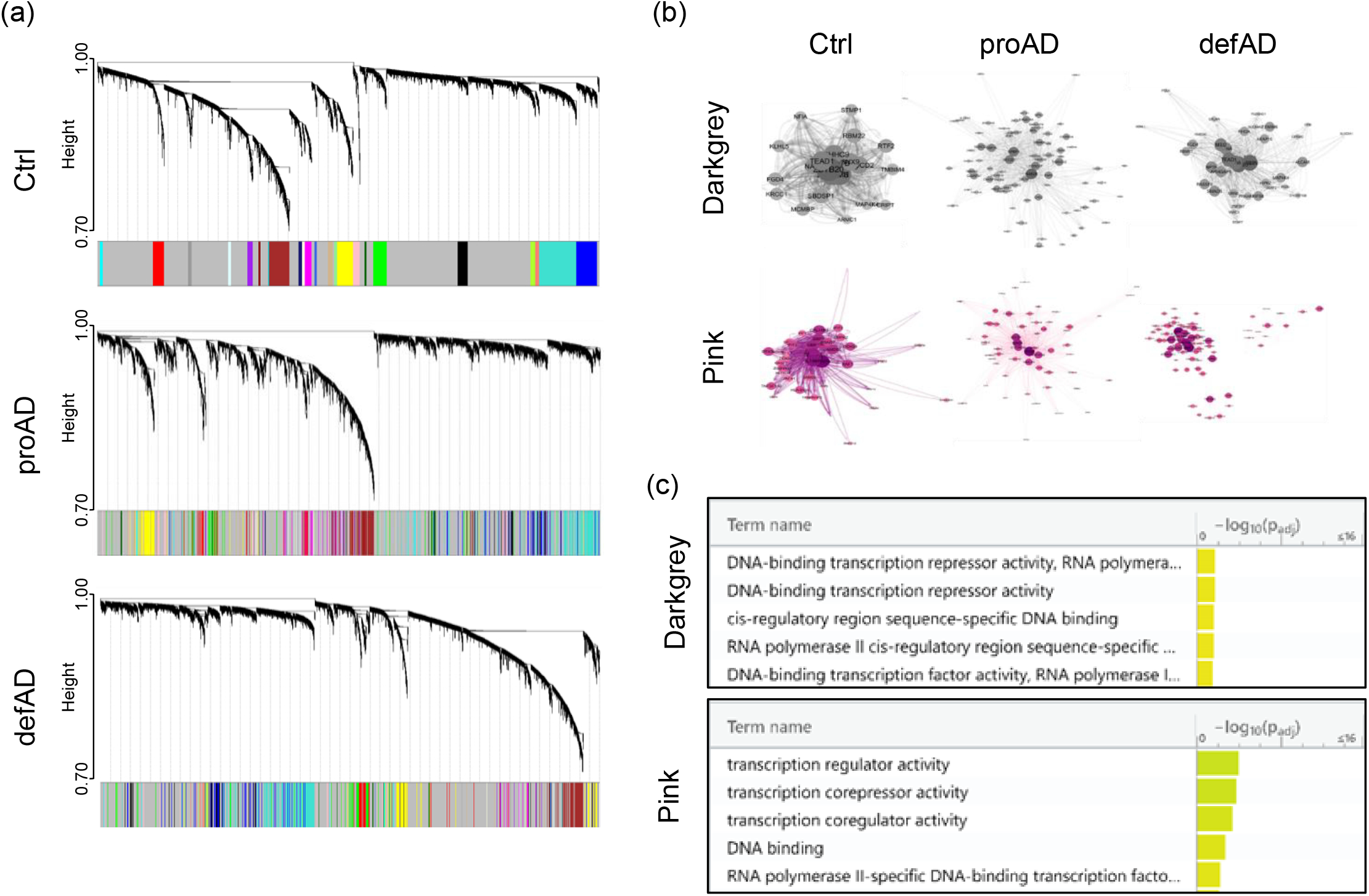
Network analysis of the AD temporal cortex shows a progressive alteration of transcriptomic pathways. (a) WGCNA coregulation analysis of control (Ctrl), proAD and defAD human temporal lobe. (b) Network representation of genes of the Darkgrey and Pink modules in Ctrl, proAD and defAD brains. (c) Transcription-related terms in the GO analysis of the Darkgrey and Pink modules. Top 5 transcription-related GO pathways have been reported.

### Tau overexpression causes global alteration of gene expression in differentiated neuroblastoma cells

The WGCNA suggested that pathways associated with the regulation of gene expression are progressively deregulated in AD. It is well known that during the progression of the disease, the amounts of total and soluble Tau protein progressively increase even before significant hyperphosphorylation or aggregation, suggesting an early imbalance of Tau cellular levels (Han et al., 2017; Kurbatskaya et al., 2016). Remarkably, we previously observed that the increase in soluble Tau leads to its accumulation in the nucleus, which alters the expression of disease-related genes in the early stages (Siano et al., 2020; Siano et al., 2019a, 2019b). Based on this evidence, we investigated whether Tau could directly modulate nuclear pathways with a particular focus on the early stages.

To accomplish this, we overexpressed Tau 0N4R, one of the most abundant Tau isoforms in the adult human brain, in differentiated neuroblastoma cells, a cellular model previously used to study molecular mechanisms in tauopathies (Bell and Zempel, 2022; Siano et al., 2023). The expression efficiency was confirmed by immunoblot analysis (Figure 2A), and global transcript expression was quantified by RNA sequencing (RNA-seq) (dataset reference number GSE239956). Principal component analysis (PCA) was applied to visualize the data and confirmed that the increased expression of Tau is the main factor contributing to variation in the global transcriptome profile (Fig 2B). RNA-seq data analysis revealed 3235 differentially expressed genes (DEGs) between Tau-overexpressing and control cells (Fig 2C). KEGG analysis identified several pathways significantly overrepresented among the DEGs, the most significant of which are listed in Table 1. Many of these terms were related to neuronal and synaptic pathways, confirming our previous observation that Tau modulates the expression of neuronal genes and might participate in triggering synaptic dysfunction occurring in early AD stages (Siano et al., 2019b).

**Figure 2.**
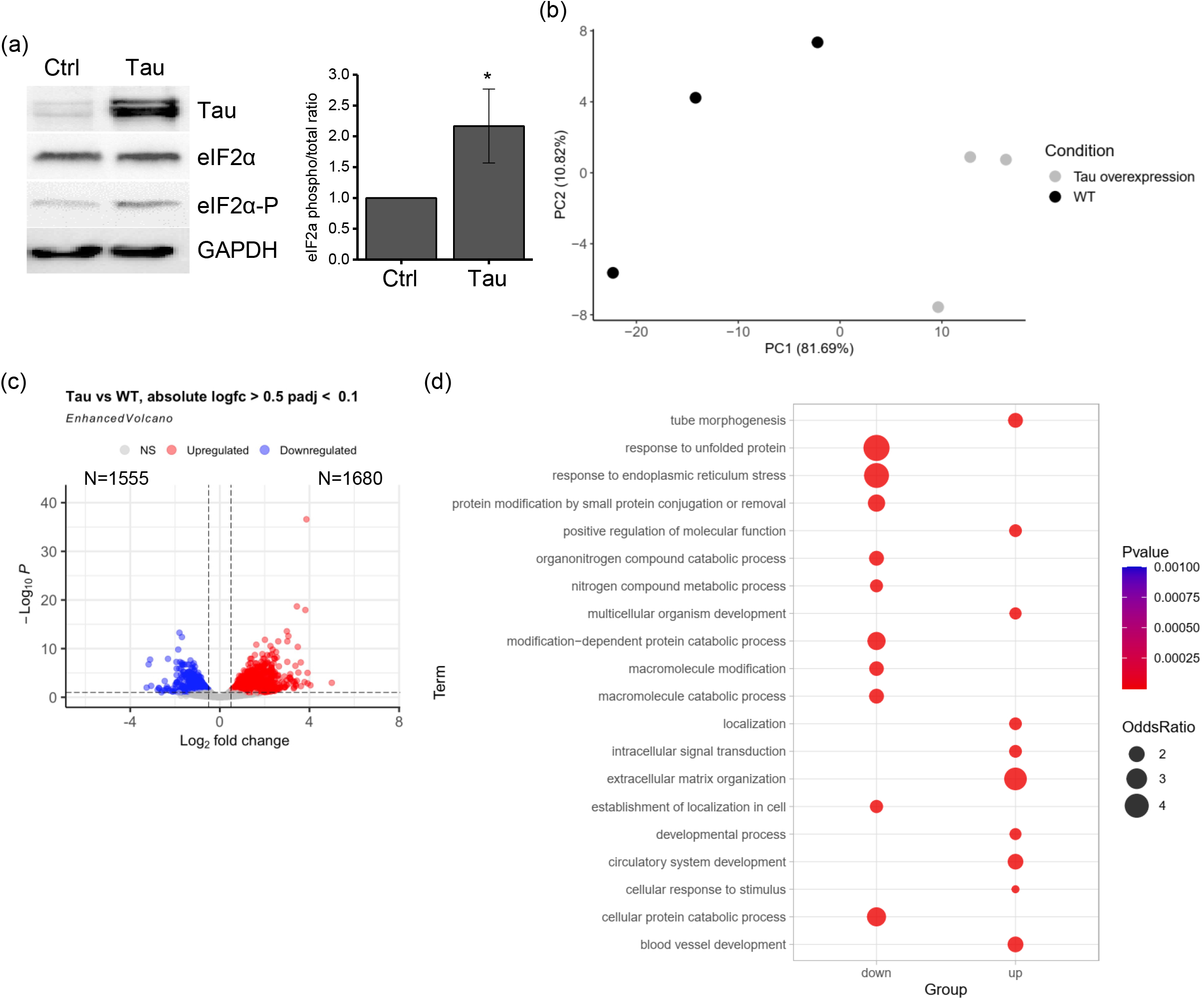
Tau overexpression in neuroblastoma cell lines leads to transcriptomic alterations. (a) WB and relative quantification of differentiated SH-SY5Y cells overexpressing Tau (proAD-mimic; Tau) and control (Ctrl) cells. Tau, eIF2α and phosphorylated eIF2α (eIF2α-P) were measured. N=4; *p<0.05. (b) PCA of proAD-mimic and control cells. (c) Volcano plot of DEGs. Red indicates upregulated genes, and blue indicates downregulated genes with |logFC| > 0.5 and FDR < 0.1. (d) GO analysis of upregulated and downregulated genes.

**Table 1.** Gene pathways altered by Tau. KEGG pathways of genes altered by Tau in neuroblastoma cell lines. Synaptic and neuronal pathways are shown in light blue.

| ID | KEGG pathway | N | Pvalue |
| --- | --- | --- | --- |
| hsa04974 | Protein digestion and absorption | 20 | 1.38E-05 |
| hsa04141 | Protein processing in endoplasmic reticulum | 53 | 2.04E-04 |
| hsa04261 | Adrenergic signaling in cardiomyocytes | 33 | 2.61E-04 |
| hsa04024 | cAMP signaling pathway | 45 | 2.82E-04 |
| hsa04512 | ECM-receptor interaction | 22 | 2.86E-04 |
| hsa04918 | Thyroid hormone synthesis | 18 | 8.21E-04 |
| hsa04925 | Aldosterone synthesis and secretion | 22 | 9.15E-04 |
| hsa04022 | cGMP-PKG signaling pathway | 28 | 1.04E-03 |
| hsa05414 | Dilated cardiomyopathy | 25 | 1.25E-03 |
| hsa04919 | Thyroid hormone signaling pathway | 19 | 1.61E-03 |
| hsa04933 | AGE-RAGE signaling pathway in diabetic complications | 25 | 2.04E-03 |
| hsa05031 | Amphetamine addiction | 18 | 2.08E-03 |
| hsa04926 | Relaxin signaling pathway | 26 | 2.55E-03 |
| hsa05206 | MicroRNAs in cancer | 40 | 2.71E-03 |
| hsa05412 | Arrhythmogenic right ventricular cardiomyopathy | 22 | 4.72E-03 |
| hsa05214 | Glioma | 14 | 5.78E-03 |
| hsa04020 | Calcium signaling pathway | 49 | 7.90E-03 |
| hsa04928 | Parathyroid hormone synthesis, secretion and action | 27 | 8.13E-03 |
| hsa04151 | PI3K-Akt signaling pathway | 66 | 9.06E-03 |
| hsa04972 | Pancreatic secretion | 21 | 9.53E-03 |
| hsa05165 | Human papillomavirus infection | 64 | 9.58E-03 |
| hsa05210 | Colorectal cancer | 19 | 9.79E-03 |
| hsa04510 | Focal adhesion | 49 | 1.24E-02 |
| hsa04730 | Long-term depression | 14 | 1.61E-02 |
| hsa04961 | Endocrine and other factor-regulated calcium reabsorption | 14 | 1.66E-02 |
| hsa04814 | Motor proteins | 36 | 1.69E-02 |
| hsa04727 | GABAergic synapse | 18 | 1.76E-02 |
| hsa04970 | Salivary secretion | 16 | 1.82E-02 |
| hsa04068 | FoxO signaling pathway | 27 | 1.85E-02 |
| hsa00564 | Glycerophospholipid metabolism | 20 | 2.01E-02 |
| hsa04360 | Axon guidance | 44 | 2.07E-02 |
| hsa05410 | Hypertrophic cardiomyopathy | 25 | 2.44E-02 |
| hsa04921 | Oxytocin signaling pathway | 31 | 2.44E-02 |
| hsa01521 | EGFR tyrosine kinase inhibitor resistance | 17 | 2.87E-02 |
| hsa04915 | Estrogen signaling pathway | 24 | 3.09E-02 |
| hsa04613 | Neutrophil extracellular trap formation | 16 | 3.14E-02 |
| hsa04218 | Cellular senescence | 31 | 3.31E-02 |
| hsa04927 | Cortisol synthesis and secretion | 15 | 3.61E-02 |
| hsa05223 | Non-small cell lung cancer | 15 | 3.69E-02 |
| hsa05166 | Human T-cell leukemia virus 1 infection | 43 | 3.76E-02 |
| hsa05146 | Amoebiasis | 16 | 4.18E-02 |
| hsa04350 | TGF-beta signaling pathway | 29 | 4.35E-02 |
| hsa04611 | Platelet activation | 20 | 4.39E-02 |
| hsa04270 | Vascular smooth muscle contraction | 27 | 4.43E-02 |
| hsa05200 | Pathways in cancer | 96 | 4.53E-02 |
| hsa00970 | Aminoacyl-tRNA biosynthesis | 11 | 4.56E-02 |
| hsa04216 | Ferroptosis | 12 | 4.59E-02 |
| hsa05034 | Alcoholism | 27 | 4.65E-02 |
| hsa04060 | Cytokine-cytokine receptor interaction | 28 | 4.79E-02 |

Upon Tau overexpression, 1680 DEGs were upregulated and 1555 were downregulated with respect to their control expression (Fig 2C). GO overrepresentation analysis of the upregulated pathways showed that extracellular matrix organization and signalling were among the most overrepresented terms, suggesting possible activation of these functions, which are increasingly investigated and are known to be relevant in tauopathies (Li et al., 2017; Ma et al., 2020; Schmidt et al., 2022) (Fig 2D; Fig EV1). For downregulated pathways, the overrepresented terms were associated with RNA regulation, protein metabolism and transport, indicating a Tau-dependent alteration of macromolecule biosynthesis (Fig 2D; Fig EV2). To functionally validate the transcriptome analysis, we assessed the phosphorylation of eIF2α (eIF2α-P) as a marker of translational inhibition induced by mechanisms involved in the response to ER stress that are altered in the cellular model (Fig 2A). Indeed, phosphorylation of eIF2α was significantly increased after Tau overexpression, supporting the link between Tau-dependent gene expression regulation and functional cellular changes.

### Tau-dependent alteration of global gene expression *in vitro* resembles early AD conditions

To assess the importance of using a neuronal cell line to investigate Tau roles, we compared our SH-SY5Y data to the transcriptome of HEK cells with inducible Tau overexpression obtained by *Montalbano et al.* (Montalbano et al., 2021). Regardless of whether they observed Tau-dependent changes in gene levels (Montalbano et al., 2021), gene expression was not correlated between the two sets (R = 0.16 and p = 0.1) (Fig EV3), thus showing that the cell model of choice has a significant impact.

We compared our RNA-seq data with DEGs in various AD stages in humans from GSE84422 to ascertain whether Tau-dependent transcriptomic alteration replicates the dysregulation of genes in particular AD phases. The proAD condition significantly overlapped our dataset (Fig 3A). We found a positive correlation between our dataset and proAD (R=0.58 and p < 2.2e-16; Fig 3B) by plotting the log fold changes (log2FCs) of our DEGs and posAD, proAD, and defAD. Approximately 75% of common genes displayed the same direction of regulation. According to this analysis, the Tau-dependent gene alterations in the proAD-mimic cellular model represented a developing intermediate AD condition in the temporal region. In addition, there was a negative correlation for posAD and no correlation for defAD (posAD: R= -0.3 and p < 0.00022; defAD: R=-0.086 and p value = 0.43; Fig 3C-D), with approximately 60-65% common genes in the second and fourth quadrants. Upon focusing on the amplitude and direction of the log2FCs of common DEGs between our dataset and proAD, we observed that while most genes showed the same direction, the amplitude was higher in the proAD-mimic model (Fig 3E), suggesting an exacerbation of the pathological phenotype in the cellular system.

**Figure 3.**
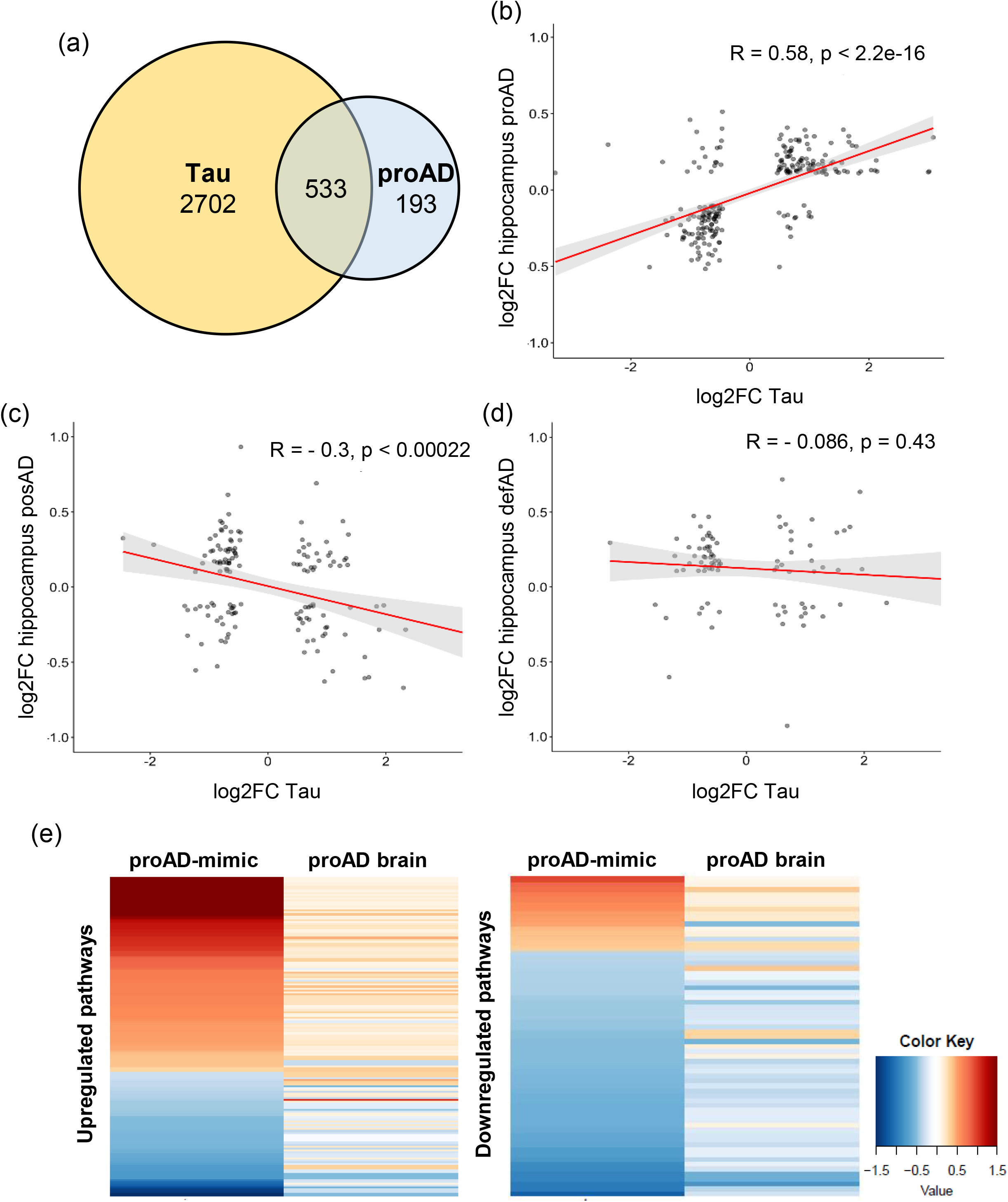
Tau-dependent transcriptome alteration correlates with the proAD stage in the human temporal region. (a) Commonly altered genes between proAD-mimic cells and proAD human brains. (b) Scatter plot representing the log2FCs of genes in proAD-mimic cells (x-axis) and proAD human brains (y-axis). (c) Scatter plot representing the log2FCs of genes in proAD-mimic cells (x-axis) and posAD human brains (y-axis). (d) Scatter plot representing the log2FCs of genes in defAD-mimic cells (x-axis) and proAD human brain (y-axis). (e) log2FC amplitude comparison between common genes in proAD-mimic cells and proAD brains.

### Tau-dependent chromatin modifications occur early in both cellular models and human brains

The GO analysis of common DEGs revealed several pathways that were differentially altered during disease progression. Terms related to transcription and chromatin modifications were uncorrelated or anticorrelated in the posAD and defAD stages, while we found a positive correlation in proAD (Figure 4A). This result suggests a key role for Tau in early pathology and shows that the increases in total and soluble Tau can lead to the alteration of transcription networks, as identified by WGCNA (Han et al., 2017; Kurbatskaya et al., 2016). Based on these data, we hypothesize that destabilized and early-aggregated Tau might alter the expression of genes related to chromatin structure and organization, thus influencing genome stability and transcription.

**Figure 4.**
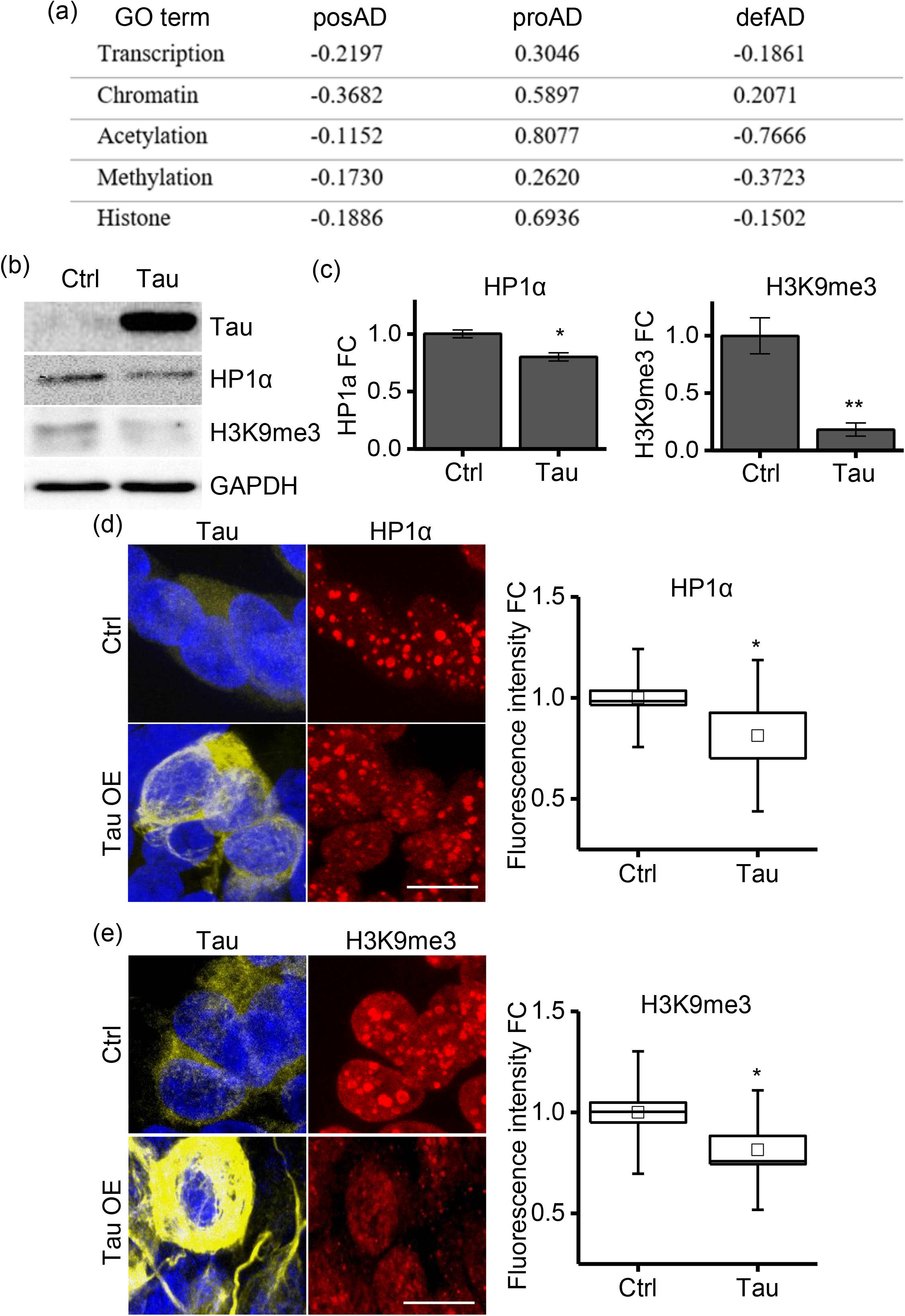
Chromatin remodelling pathways are altered by Tau. (a) Correlation analysis of DEGs in chromatin remodelling pathways in posAD, proAD and defAD. (b) WB of the heterochromatin markers HP1α and H3K9me3 in proAD-mimic (Tau) and control (Ctrl) cells. (c) Bar plot of HP1α and H3K9me3 in proAD-mimic (Tau) and control (Ctrl) cells. (d) Immunofluorescence and relative quantification of HP1α in proAD-mimic and control cells. Red: HP1α; yellow: Tau; blue: DAPI. (e) Immunofluorescence and relative quantification of H3K9me3 in proAD-mimic and control cells. Red: H3K9me3; yellow: Tau; blue: DAPI. Scale bar: 5 µm *p<0.05; **p<0.01.

To elucidate chromatin remodelling deregulation, we detected the heterochromatin markers HP1α and H3K9me3, and we observed a significant reduction in the proAD-mimic model cells compared to control cells by both immunoblotting (WB) (Fig 4B-C) and immunofluorescence (IF) (Fig 4D-E). These results indicate that increased Tau levels alter the level of chromatin remodelling factors, leading to a reduction in heterochromatin that mimics an early/intermediate AD condition.

To confirm these results, we analysed the temporal lobes of control (BS1/2) and AD patients at early/intermediate pathological stages. By WB analysis, we observed a significant reduction in HP1α but not H3K9me3 levels (Fig 5A) at Braak stages 3 and 4 (BS3/4) compared to those in controls (BS1/2). In IF experiments, we labelled AD temporal lobes with the Tau^AT8^ phospho-epitope as a marker of Tau pathology and NeuN as a marker of neuronal cells. At BS3 and BS4, we observed a sparse AT8 signal in a small percentage of NeuN^+^ cells. We analysed the heterochromatin marker H3K9me3, and we observed that the total fluorescence was not altered as much as in the WB experiment. However, upon measuring the H3K9me3 signal in Tau^AT8^-positive neurons, we observed a significantly weaker signal than that in healthy brains (Fig 5B). Remarkably, at late AD stages (BS5/6), the AT8 signal was detected in all tissues as expected, and the total H3K9me3 fluorescence was significantly reduced, further supporting our evidence regarding the Tau-dependent alteration of heterochromatin (Fig 5A-B). This finding further supports the interplay between pathological Tau and heterochromatin changes in stages that anticipate the defAD condition.

**Figure 5.**
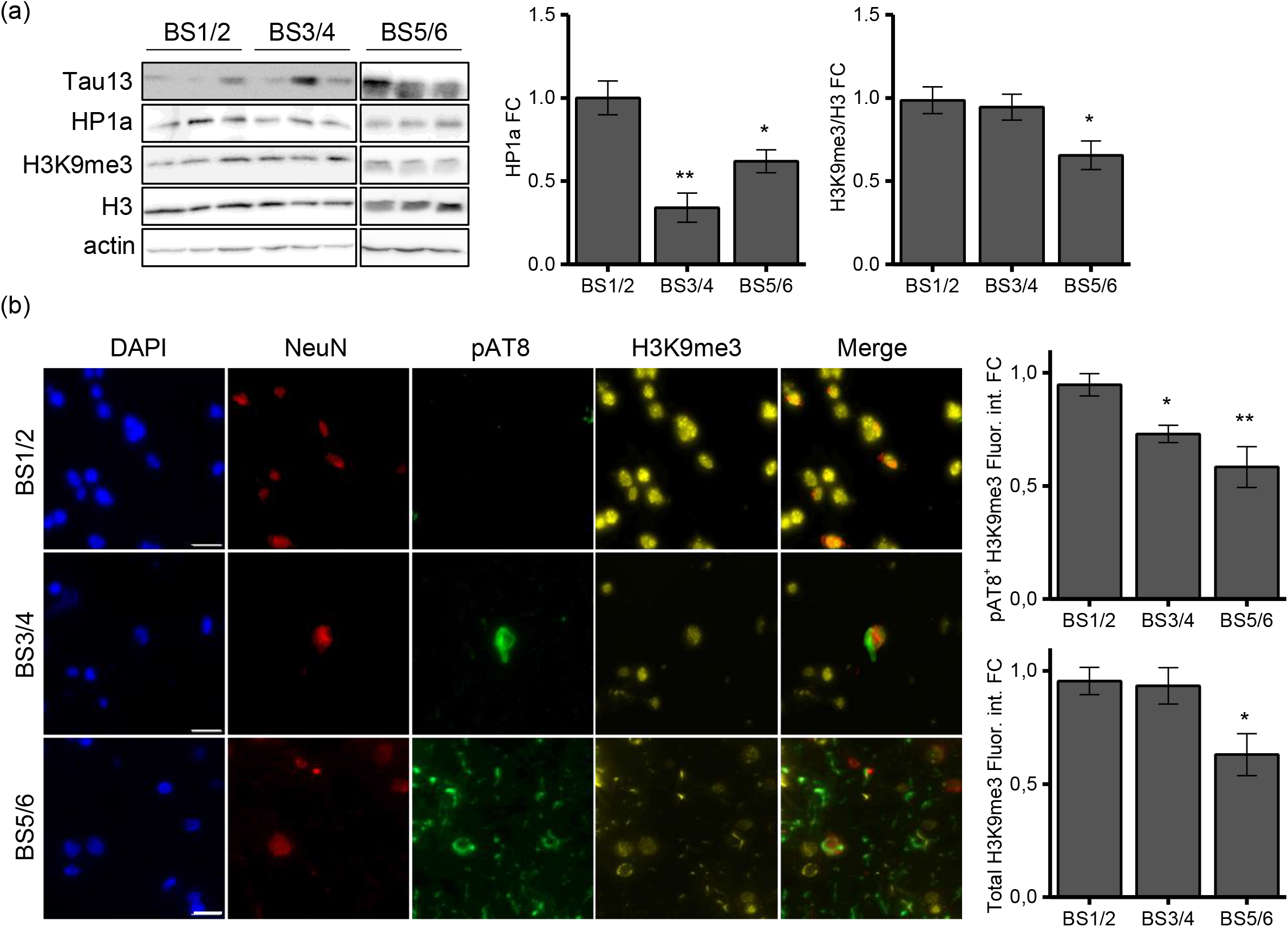
Heterochromatin changes are associated with Tau pathology in AD temporal lobes at intermediate stages. (a) WB of heterochromatin markers HP1α and H3K9me3 in human brain samples at BS1/2 (N=7), BS3/4 (N=6) and BS5/6 (N=8) and relative quantification. (b) Immunofluorescence and relative quantification of H3K9me3 in human brain samples at BS1/2 (N=6), BS3/4 (N=5) and BS5/6 (N=5). Red: NeuN; yellow: H3K9me3; green: pAT8; blue: DAPI. Scale bar: 20 µm *p<0.05; **p<0.01.

## Discussion

The Tau protein has been extensively studied for its ability to bind and stabilize MTs and to undergo posttranslational modifications that alter its affinity to tubulin, leading to cytoskeletal instability and the formation of toxic amyloidogenic aggregates (Wang and Mandelkow, 2016). It is generally assumed that these key events mediate the neurodegenerative process; however, these observations cannot entirely explain the sequential cellular alterations occurring during the progression of tauopathy. We recently demonstrated that Tau overexpression or destabilization results in significant increases in both the cytoplasmic and nuclear pools of Tau, thus mimicking an early pathological condition. We have described a pathological stage-dependent role of Tau in the nuclear compartment associated with alterations in glutamatergic gene expression and downstream neuronal transmission (G. Siano et al., 2020; Siano et al., 2019b).

By using unbiased WGCNA of microarray data from human temporal lobes at different stages (posAD, proAD and defAD), we observed a progressive loss of coregulation of genes associated with AD pathways such as synaptic transmission and neuronal function, as already observed (Hu et al., 2020; Miller et al., 2008; Sun et al., 2022). Of note, we also identified pathways related to chromatin and transcription, suggesting that these mechanisms can undergo functional changes during the course of the disease. Transcription-associated modules have not been described in previous coregulation analyses employing WGCNA on AD samples. Moreover, in longitudinal studies, either the number of samples has been low, thus limiting the depth of the analysis, or the tissues considered have not been brain-derived. In any case, the previous studies enabled us to identify not only relevant pathways associated with AD pathology but also gene candidates that can drive the pathology or be employed as biomarkers (Hu et al., 2020; Miller et al., 2008; Soleimani Zakeri et al., 2020; Sun et al., 2022; Upadhyaya et al., 2020; Zhang and Kiryu, 2023). Here, the dataset employed provided transcriptomic data of early, intermediate and late AD stages identified via integration of several pathological parameters. Moreover, the higher number of subjects per group compared with those in other WGCNAs increased the sensitivity of the coregulation network, thus revealing pathways not previously described, such as those related to chromatin and transcription. Given our previous observations on the subcellular imbalance of Tau in early stages and its involvement in gene expression modulation, we focused on the association between Tau and global transcriptional changes.

By RNA-seq analysis of differentiated SH-SY5Y cells transfected with the wild-type Tau 0N4R isoform, we observed a great change in global gene expression, with more than 3000 genes significantly altered. This evidence suggests that the increased availability of Tau leads to an alteration in synaptic genes as well as a global transcriptomic rearrangement that might affect cellular physiology (Siano et al., 2021). Our findings are further supported by a similar study in human embryonic kidney (HEK) cells in which the authors observed a Tau-dependent transcriptomic change that impacted several functions. Although the cellular model was unrelated to neurons and the transcriptomic technique employed has some limitations compared to RNA-seq (Elrod et al., 2019; Montalbano et al., 2021), the study confirmed that Tau-dependent gene modulation is an intrinsic function of the Tau protein and is independent of the cellular background. However, the low correlation between our dataset and the dataset from *Montalbano et al.* indicates that the outcome of research on the role of Tau in gene expression significantly relies on the cellular model employed.

A transcriptome shift is generally associated with functional alterations leading to pathological conditions. The GO analysis identified terms related to synaptic functions as Tau-dependent pathways, as expected (G. Siano et al., 2020; Siano et al., 2019b). Additionally, we found terms that are relevant for neuronal physiology and pathology, such as protein metabolism, which is well known to have a key role in tauopathy progression, and less canonical pathways, such as those related to RNA metabolism or extracellular matrix remodelling. These proteins are known to be altered in tauopathies but are still not completely characterized (Chauderlier et al., 2018; Hernández-Ortega et al., 2016; Ma et al., 2020; Violet et al., 2014). To functionally validate our transcriptomic analysis, we measured the phosphorylation of eIF2α in our cellular model and observed a significant increase in eIF2α-P after Tau overexpression. Phosphorylation of eIF2α is a well-known marker of response to ER stress and translational mechanisms in tauopathies (Clemens, 2001; Gerakis and Hetz, 2018; Radford et al., 2015; Shacham et al., 2021). Our findings indicate that these pathways, which are essential for neuronal functionality, are transcriptionally altered by Tau.

The data collected with the cellular model, here defined as proAD-mimic, showed that changes in Tau cellular availability affected the global transcriptome. We performed a meta-analysis to compare the transcriptome of this model with the datasets from temporal regions at three AD stages: posAD, proAD and defAD. The meta-analysis revealed a significant overlap of common DEGs specifically with the proAD stage, supporting the conclusion that the gene expression changes caused by Tau reflect a pathological condition that precedes the late AD phase (Siano et al., 2020). A gene-by-gene FC comparison further indicated a significant positive correlation between the proAD-mimic and proAD brains, supporting the hypothesis that the transcriptomic modifications observed in patients are also mediated by Tau pathology, as previously suggested (Siano et al., 2021). Indeed, in proAD conditions, the Tau protein is destabilized, oligomerized, and highly soluble (Alonso et al., 1997; Guo et al., 2017; Lowe et al., 2018). Furthermore, the total and soluble amounts of Tau are significantly elevated in intermediate AD stages, especially in frontal and temporal human brain regions, before Tau aggregates form. This suggests that the increase in total Tau level can impair neuronal homeostasis and drive the pathological changes observed in late phases (Han et al., 2017; Kurbatskaya et al., 2016). In this context, several neuronal functions are compromised, such as synaptic transmission and DNA repair, and our findings suggest that these pathways could be transcriptionally altered by Tau (Colnaghi et al., 2020; Siano et al., 2021). In addition, the meta-analysis identified the proAD mimic as a reliable and easy model to resemble a developing intermediate stage (proAD) *in vitro*. Remarkably, in the proAD mimic, the FC amplitude of DEGs was higher than that in proAD, suggesting that in this model, the pathological phenotype is exacerbated due to the high percentage of Tau-overexpressing cells or due to the higher amount of Tau. This worsening might be weakened in the brain by the cellular heterogeneity of the tissue, which could dampen the global impact of neuronal transcriptomic changes.

The evidence regarding the specificity of the proAD-mimic model in resembling proAD tissue was further strengthened by the comparison with posAD and defAD. The posAD and defAD transcriptomes were negatively correlated and not correlated with our dataset, respectively. The negative correlation with posAD might suggest early transcriptomic compensation to prevent Tau-dependent alteration of gene expression and to maintain neuronal homeostasis, since at this early stage, Tau undergoes initial destabilization and oligomerization. In contrast, the loss of correlation with the terminal stages could be the consequence of excessive transcriptomic and functional damage in proAD leading to global gene dysregulation. This evidence closely relates to our previous study on the human prefrontal cortex at different Braak stages. We observed significant deregulation of synaptic genes mainly at Braak stages 3 and 4 (corresponding to an early or intermediate AD stage), and we hypothesized a causal link with destabilized and oligomerized Tau protein (G. Siano et al., 2020). Our meta-analysis further supports this thesis, confirming that in early and intermediate pathological stages, there is a Tau-dependent transcriptomic alteration impinging on the synaptic pathway.

The meta-analysis revealed that Tau has a significant role in the transcriptomic modifications observed in proAD and indicated the Tau-dependent pathways altered in proAD and how they are regulated in the previous and following phases. The GO analysis of common genes between our dataset and proAD brains identified several terms related to transcription and chromatin remodelling, according to the WGCNA data. The correlation analysis of epigenetic terms showed a positive correlation only with proAD, as expected, whereas a negative correlation and no regulation were observed in the posAD and defAD stages, respectively. The Tau-dependent alteration of chromatin remodelling pathways associated with the proAD stage indicates an early modification of chromatin structures (Eissenberg and Elgin, 2014; El Hajjar et al., 2019; Frost et al., 2014; Napoletano et al., 2021). The reductions in the levels of heterochromatin markers, HP1α and H3K9me3, in the proAD-mimic model indicated Tau as a major factor leading to the loss of heterochromatin. This evidence was further supported by the analysis of the same markers in human temporal tissues, in which at early/intermediate stages, a reduction in the H3K9me3 signal was observed specifically in neurons that exhibited hyperphosphorylation of Tau at the AT8 epitope, a site commonly associated with Tau pathology (Neddens et al., 2018). This result validated our previous observations, indicating that heterochromatin loss is strictly connected with Tau. However, HP1α showed a stronger and wider reduction than H3K9me3. It is conceivable that alteration of HP1α, a general marker of heterochromatin and the nucleolus, could be more responsive to minor Tau perturbations or that another mechanism could have contributed to its reduction. These data imply that Tau can affect chromatin pathways and cause heterochromatin loss in intermediate AD stages, thus influencing transcriptional mechanisms.

We previously demonstrated that increased Tau availability causes an imbalance in Tau amounts in different subcellular compartments, mimicking an early pathological condition. We observed that this imbalance, particularly in the nuclear compartment, causes alterations in glutamatergic genes. (Siano et al., 2021, 2019b). Here, we found that Tau alters not only glutamatergic pathways but also other neuronal pathways associated with stages preceding severe AD pathology that are characterized by higher Tau levels in turn (Han et al., 2017; Kurbatskaya et al., 2016). In fact, we observed early alterations of chromatin remodelling genes that could affect chromatin structure, thus enhancing transcriptional deregulation with a positive feedback mechanism. A connection between Tau and chromatin structure has been previously described since Tau directly interacts with TRIM28, a key protein involved in histone modifications. Indeed, we recently demonstrated that Tau can cause the delocalization of the chromatin remodeller HDAC1, an event associated with the reduction of heterochromatin (Lagger et al., 2002; Siano et al., 2023), suggesting one of the possible mechanisms mediating the Tau-dependent epigenetic and transcriptional alteration. Moreover, Tau activates transposons in silent regions, leading to genomic instability, even though this event is associated with neurodegeneration (Guo et al., 2018; Rousseaux et al., 2016; Sun et al., 2018). Notably, Tau binds directly to DNA with a preference for specific sequences and histones, thus contributing to chromatin organization (Benhelli-Mokrani et al., 2018; Mansuroglu et al., 2016; Rico et al., 2021). All these studies indicate that Tau has a prominent role in affecting chromatin structure, and here, we showed the involvement of Tau in phases that anticipate the defAD condition. Moreover, our results suggest that chromatin pathway changes are downstream consequences of altered Tau availability.

Recent studies have reported that the progression of neurodegeneration in the prefrontal cortex of AD brains is the result of a multi-scale disruption of genome organization. This evidence indicates that the chromatin landscapes are dynamically altered, causing the deregulation of transcriptomic pathways, resulting in cell-type specific dysfunction. Furthermore, genome instability has been found to modulate pathways related to synaptic signalling, neurogenesis, cell adhesion, immunity, cell signalling, and RNA metabolism (Dileep et al., 2023; Gazestani et al., 2023; Xiong et al., 2023). It is noteworthy that in our system, the modulation of Tau leads to a similar reorganization, suggesting that Tau dysfunction might be hierarchically upstream of these mechanisms.

Chromatin remodelling is closely related to transcriptome alterations, and we hypothesize that the progressive increase in Tau availability causes early perturbation of upstream regulators of gene expression via direct interaction with chromatin, chromatin remodellers, or transcription factors or indirectly via modification of cofactor function. This upstream perturbation could lead to further changes in chromatin structure and consequently to a global transcriptomic alteration that feeds into a self-perpetuating mechanism, causing a progressive disruption of transcriptome homeostasis and neuronal functions, which drives neurodegeneration. Remarkably, Tau-dependent alteration of epigenetic genes has also been recently observed in nonneuronal cells, supporting our observation in a neuronal background (Montalbano et al., 2021). The links between those modifications and the early stages of AD from our data suggest that neuronal damage caused by pathological Tau occurs and must be prevented early by targeting cellular mechanisms associated with transcription regulation.

## Experimental Procedures

### Cell culture, transfection and differentiation

SH-SY5Y neuroblastoma cells were maintained in DMEM/F12 (Gibco) supplemented with foetal bovine serum (FBS) and penicillin/streptomycin. For the transcriptomic experiments, 2 x 10^5^ cells were plated in p30 wells and transfected the day after seeding by Lipofectamine 2000 according to the manufacturer’s instructions. The day after transfection, cells were differentiated by 10 µM retinoic acid (Sigma‒Aldrich) for 5 days followed by 50 ng/ml BDNF (Alomone Labs) for 15 days in serum-free DMEM/F12 medium. After differentiation, cells were collected for Western blotting and RNA extraction. For Tau overexpression, the cDNA encoding Tau isoform D (383 aa) was cloned into the BglII site of pcDNA3.1, and an empty pcDNA3.1 vector was used as a control as previously described (Siano et al., 2019b).

### WB analysis

For WB analysis, experiments were performed as previously described (Giacomo Siano et al., 2020). In brief, total protein extracts were prepared in lysis buffer supplemented with protease and phosphatase inhibitors (Roche). Proteins were quantified by bicinchoninic acid (BCA) assay (Thermo Fisher Scientific). Twenty micrograms of total protein were loaded for each sample. The proteins were separated by SDS–PAGE and electroblotted onto Hybond-C-Extra (Amersham Biosciences) nitrocellulose membranes. The membranes were blocked with 5% skimmed milk powder in TBS and 0.1% Tween 20. The antibodies for WB were as follows: mouse anti-Tau (Tau13) 1:1000 (Santa Cruz Biotechnology), rabbit anti-HP1α 1:500 (Abcam), rabbit anti-H3K9me3 1:500 (Abcam), rabbit anti-eIF1α 1:1000 (Cell Signaling), rabbit anti phospho-eIF1α 1:1000 (Cell Signaling), mouse anti-GAPDH 1:10000 (Santa Cruz Biotechnology), HRP-conjugated anti-mouse 1:1000 (Santa Cruz Biotechnology), and HRP-conjugated anti-rabbit 1:1000 (Santa Cruz Biotechnology). Western blot quantification was performed using ImageJ software.

### Immunofluorescence

For IF experiments *in vitro*, cells were fixed with ice-cold 100% methanol for 5 min. The cells were permeabilized (PBS, 0.1% Triton X-100), blocked (1% wt/vol BSA) and incubated with primary and secondary antibodies. The slides were mounted with VECTASHIELD mounting medium (Vector Laboratories). The antibodies for IF were as follows: mouse anti-Tau (Tau13) 1:500 (Santa Cruz Biotechnology), rabbit anti-HP1a 1:250 (Abcam), rabbit anti-H3K9me3 1:500 (Abcam), mouse anti-Tau AT8 1:250 (Invitrogen), anti-NeuN (Boster Biological Technology) 1: 300, anti-mouse Alexa Fluor 633, and anti-rabbit Alexa Fluor 488 (Life Technologies). Nuclear staining was performed by incubation for 15 min with DAPI. Images were acquired on a Zeiss Laser Scanning (LSM) 880 confocal microscope (Carl Zeiss, Jena, Germany) supplied with GaAsP (gallium:arsenide:phosphide) detectors. The samples were viewed with a 63X Apochromat oil immersion (1.4 NA) DIC objective. Whole-cell images were acquired with a z-stack series with 0.5 µm intervals and summed with the z-projection tool from Fiji. Chromatin marker fluorescence was analysed by Fiji software.

Freshly frozen brain tissue was sectioned at 10 μm using a cryostat (Thermo Fisher Scientific). The brain slices were fixed in ice-cold acetone for 5 minutes before being rinsed twice with PBS and twice with PBS-T and blocked in animal-free blocker for 30 minutes. The tissue sections were treated for 1 hour at room temperature with primary antibodies diluted in blocking solution, rinsed three times with PBS-T, and then incubated with secondary antibodies for 1 hour at room temperature. Following a PBS-T wash, the slices were incubated with PureBlu DAPI (Bio-Rad, Cat# 1351303) for 3 minutes before being mounted with ProLong Gold antifade mounting media (Thermo Fisher Scientific, Cat# P36934). Images were scanned using an Olympus SLIDEVIEW VS200 slide scanner.

ImageJ software was used to analyse immunofluorescence images. Image5D followed by Z Projection was employed for z-stack images. H3K9me3 fluorescence was measured in NeuN and AT8 positive cells after background subtraction. In control BS1 and BS2 samples, AT8 signal was undetectable or very low. In AD samples from BS3 to BS6 we detected a significant and increasing AT8 signal. H3K9me3 fluorescence in NeuN/AT8-positive cells was compared to NeuN-positive cells in BS1/2 samples. An average of 55 cells were analysed for each brain sample. For imaging analysis, images have not been manipulated. For representative images in the text, Brightness and Contrast plugin was used homogeneously for the different experimental groups to clarify measured differences.

### Statistical analysis

For WB and IF analyses, the non-parametric Kruskal-Wallis test was used. All results are shown as the mean ± standard error of the mean (SEM). For WB experiments, N ≥ 3 independent biological experiments were performed. For IF experiments on neuroblastoma cells: N ≥ 12; each biological replicate corresponded to one cell. For human brain sample analyses, N ≥ 4. Significance is indicated as * for p < 0.05, ** for p < 0.01, *** for p < 0.001 and **** for p < 0.0001.

### WGCNA

Public transcriptomic datasets were sourced from the Gene Expression Omnibus (GEO) platform hosted on NCBI (https://www.ncbi.nlm.nih.gov/geo/). The GSE84422 dataset (Wang et al., 2016), which includes data from 1053 postmortem brain samples across 19 brain regions from a total of 125 patients dying at different AD stages, with 50-60 subjects per brain region, was used. The patients were clustered by AD severity into 4 groups, the normal (n=110), posAD (n=110), proAD (n=83) and defAD (n=163) groups, according to their clinical dementia rating (CDR), Braak NFT score, sum of NFT density across different brain regions, average neuritic plaque density and average Consortium to Establish a Registry for Alzheimer’s Disease (CERAD) score. We considered for our analyses the CDR since it integrates several parameters to define the AD condition. Gene expression profiling was performed on the Affymetrix Human Genome U133B Array (GPL97) (Wang et al., 2016). Overrepresentation analysis (ORA) provided a knowledge-based classification of genes and list of associated pathways. For further insight into the biological functions of the sets of DEGs, without any a priori assumption on the involvement of genes in specific pathways, WGCNA was performed on a larger patient set from the GSE84422 dataset. All the samples from the temporal cortex of GSE84422 were grouped and divided into control, proAD and defAD groups. Because posAD modules were conserved when compared with the modules healthy control brains, the posAD group was excluded from the analysis. We used the WGCNA package and customized the WGCNA tutorial available at https://horvath.genetics.ucla.edu/html/CoexpressionNetwork/Rpackages and the code for building consensus networks from “Meta-analyses of data from two (or more) microarray datasets” by Jeremy Miller. The dataset was divided into subsets according to the AD stage, an appropriate soft thresholding power β (β = 9) was chosen so that the data of both subsets would fit a scale-free distribution, and the compatibility of the subsets was verified by calculating the correlation between the ranked gene expression and the correlation between the ranked connectivity. The signed adjacency matrix provided the distance measure for building modules on the control set. To detect even small gene clusters, deepSplit was set to 3, and the minimum cluster size was set to 30-3*deepSplit. Module eigengenes, gene module membership (i.e., the correlation of module genes with eigengenes) and module preservation in the disease phases were later assessed. Hub genes were defined as those with the highest module membership. Modules were visualized with Cytoscape. To choose the appropriate number of clusters, the modules were first inspected both visually and with assignment indicators with the R packages “ppclust”, “factoextra”, “cluster” and “fclust”, and then the modified partition coefficient and the partition entropy were maximized. To associate each cluster with a molecular function, gProfiler software was used, including all known genes in the GO database. Top pathways with Benjamini‒Hochberg FDR values < 0.05 defined cluster identity.

### RNA extraction and RNA-seq

Two experimental groups were compared: the control and Tau^WT^ overexpression groups. RNA was extracted with a NucleoSpin (Macherey-Nagel) RNA extraction kit according to the manufacturer’s instructions. Sequencing of RNA samples was performed using Illumina’s next-generation sequencing methodology (Bentley et al., 2008). In detail, total RNA was quantified and quality checked using Bioanalyzer 2100 instrument in combination with RNA 6000 nano kit (both Agilent Technologies). Libraries were prepared from 800 ng of input material (total RNA) using NEBNext Ultra II Directional RNA Library Preparation Kit in combination with NEBNext Poly(A) mRNA Magnetic Isolation Module and NEBNext Multiplex Oligos for Illumina (Index Primers Set 1/2/3/4) following the manufacturer’s instructions (New England Biolabs). Quantification and quality checked of libraries was done using an Bioanalyzer 2100 instrument and DNA 7500 kit (Agilent Technologies). Libraries were pooled and sequenced in two lanes on a HiSeq 2500. System runs in 51 cycle/single-end/high-throuput (SBS reagent v3) mode. Sequence information was converted to FASTQ format using bcl2fastq v1.8.4.

### RNA sequencing data analysis

The reads were mapped onto the genome (Ensembl: GRCh38.92, (Zerbino et al., 2018)) with Tophat v2.1 (Kim et al., 2013; Langmead et al., 2009) using parameter `--no-convert-bam -- no-coverage-search -x 1 -g 1`: on average, 80% of all reads were mapped univocally. The reads per gene was counted using featureCounts v1.5.0 (Liao et al., 2019). Read counts were introduced in the the statistical environment R (v.3.4.1) for further processing. Read count were normalized for reads per samples (RPM) and transcript length (RPKM) (Mortazavi et al., 2008). DEGs were defined by an FDR cut-off of < 0.1 in the statistical tests of all three R packages, edgeR (Robinson et al., 2010), DESeq (Anders and Huber, 2010) and baySeq (Hardcastle and Kelly, 2010), and by an absolute log2FC > 0.5. The 300 most variant genes were selected for the PCA plot using the R package pcaExplorer after variance stabilization (Marini and Binder, 2019). The EnhancedVolcano package was used for the volcano plot (Blighe et al., 2018).

GO analysis of the DEGs was performed by ORA with the GO Biological Process annotations from the GO.db package (Carlson, 2019). ORA was performed by a right-sided hypergeometric test, and terms with BH-adjusted p values < 0.05 were considered significant. WebGestalt was used to summarize GO entries with the affinity propagation method (Wang and Reddy, 2017). For KEGG pathway enrichment, pathways were retrieved using the KEGGREST package (Tenebaum and Maintainer, 2023), and the Wilcoxon test was performed for each pathway on the merged set of up- and downregulated DEGs. Pathways with a P value < 0.05 and number of genes > 10 are reported in the table.

### Gene expression comparison with the human dataset

For transcriptome meta-analysis, public transcriptomic datasets were sourced from the GEO platform hosted on NCBI (https://www.ncbi.nlm.nih.gov/geo/). GSE84422 was used. The patients included in the GSE84422 dataset were classified by AD stage into posAD, proAD and defAD. The R packages “GEOquery” (Davis and Meltzer, 2007), “Rcpp” (Eddelbuettel and François, 2011) and “tidyselect” were used to import datasets and create subsets by diagnosis and sample type or brain region. To compare different platforms, platform-specific IDs were converted into Entrez Gene IDs using the corresponding R annotation databases from Bioconductor, available via bioMart (version 3.8) (Durinck et al., 2009). Multiple platform IDs matching the same gene were summarized in a single entry with gene expression as the mean of probe expression. Probes mapping on the same gene with a discordant logFC were excluded from the analysis. Differential expression analysis of microarray data was performed with the R package “limma” (version 3.8) (Ritchie et al., 2015). An FDR-adjusted p value < 0.1 was set as the cut-off for differential expression. The temporal region data from the GSE84422 dataset were used. The GSE84422 dataset was compared to our SH-SY5Y data to identify the stage of the disease that was best recapitulated in the cellular model. To assess the conservation of specific pathways, genes belonging to the pathway were ranked according to their expression level in each dataset, as in Gene Set Enrichment Analysis. In this way, we could underscore the combinations of small contributions from the single genes to the whole pathway. The DEGs in our dataset (FDR < 0.1) were compared to the genes in the temporal region dataset, and the correlation of the most significant hits was computed. Genes with absolute logFCs < 0.1 were excluded from the analysis.

Enrichment of GO terms in the SH-SY5Y transcriptome was assessed by ORA as described above in “RNA sequencing data analysis”. The direction and magnitude of the fold changes of genes in the most differentially represented pathways were compared to those for patients in dataset GSE84422 to assess correlations and the ratios of fold change amplitude. All DEGs in the SH-SY5Y transcriptome were included in the analysis.

### Gene expression comparison with the iHEKTau dataset

To assess the impact of the cell line on the transcriptome after Tau overexpression, we compared our SH-SY5Y data to the transcriptome data of HEK cells with inducible Tau overexpression obtained by Montalbano *et al*. The raw data for the control cells and for iHEKTau cells were downloaded from NCBI under SRA reference number PRJNA744518 and processed as indicated in the original paper (Montalbano et al., 2021).

Genes with absolute logFC values > 0.1 and adjusted p values < 0.1 in both datasets were used to compute the correlations between gene expression levels in Tau-overexpressing SH-SH5Y and iHEKTau cells.

## Acknowledgements

The authors are grateful to M. Calvello, V. Liverani and A. Viegi for technical support, the Next Generation Sequencing facility and the Life Science Computing facility of the Leibniz Institute on Aging –Fritz Lipmann Institute-for RNA-seq and primary data analysis. We thank the Roche postdoctoral fellowship program for funding C.W. and M.T.

## Author Contributions

GS, MV, AC, CDP and AC designed the experiments, and GS, MV and CDP wrote the manuscript. GS, MV, MT and CW performed the experiments; GS, MV, AC, MG collected and analysed the data. AC provided reagents and contributed to the discussion and correction of the manuscript. MCG and JJMH contributed to the discussion and correction of the manuscript.

## Data availability

RNA-seq data of SH-SY5Y overexpressing Tau and control cells are available in NCBI-GEO platform with the reference number GSE239956 (https://www.ncbi.nlm.nih.gov/geo/query/acc.cgi?acc=GSE239956)

## Expanded View Figure legends

**Fig EV1.**
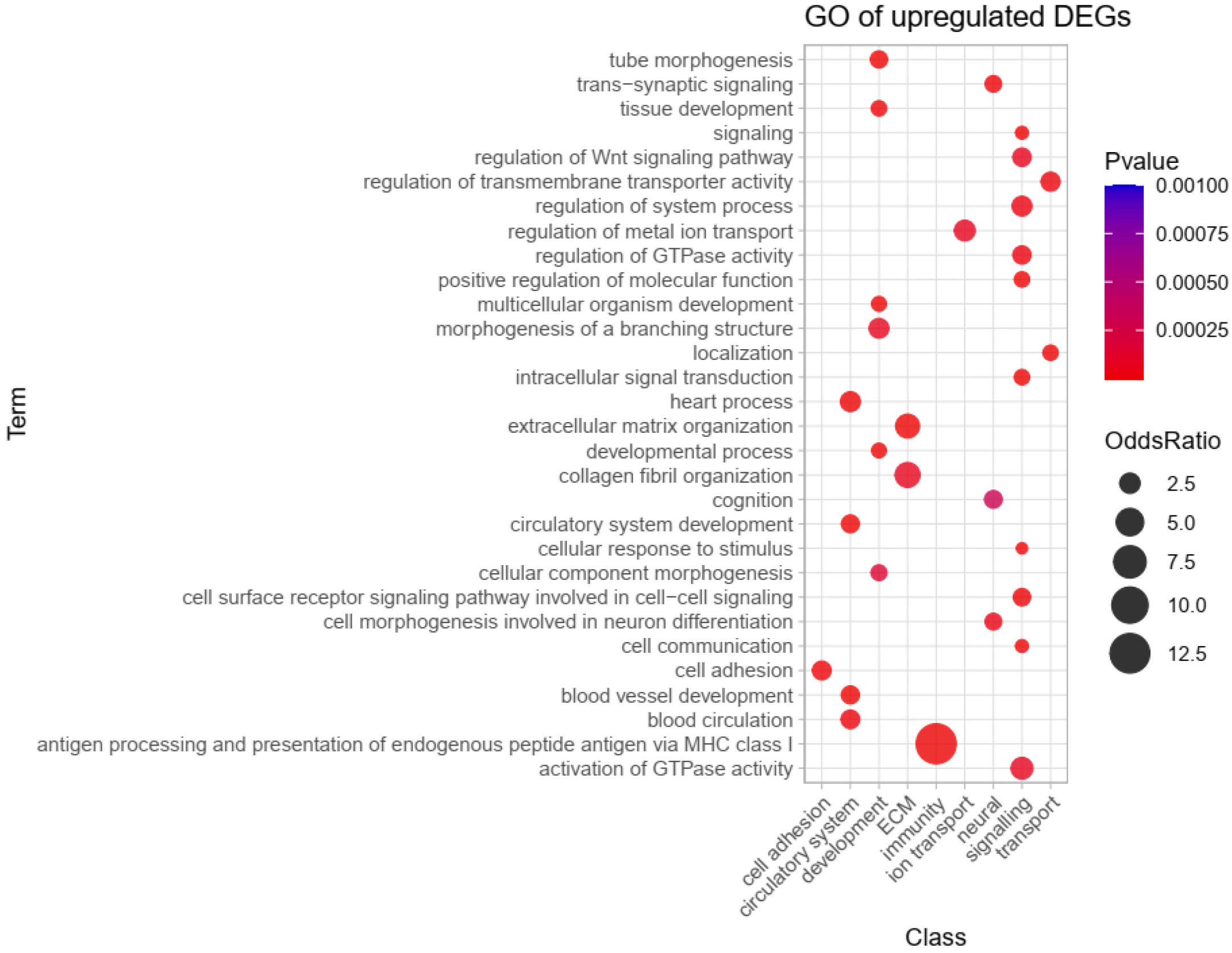
GO overrepresentation analysis of upregulated pathways.

**Fig EV2.**
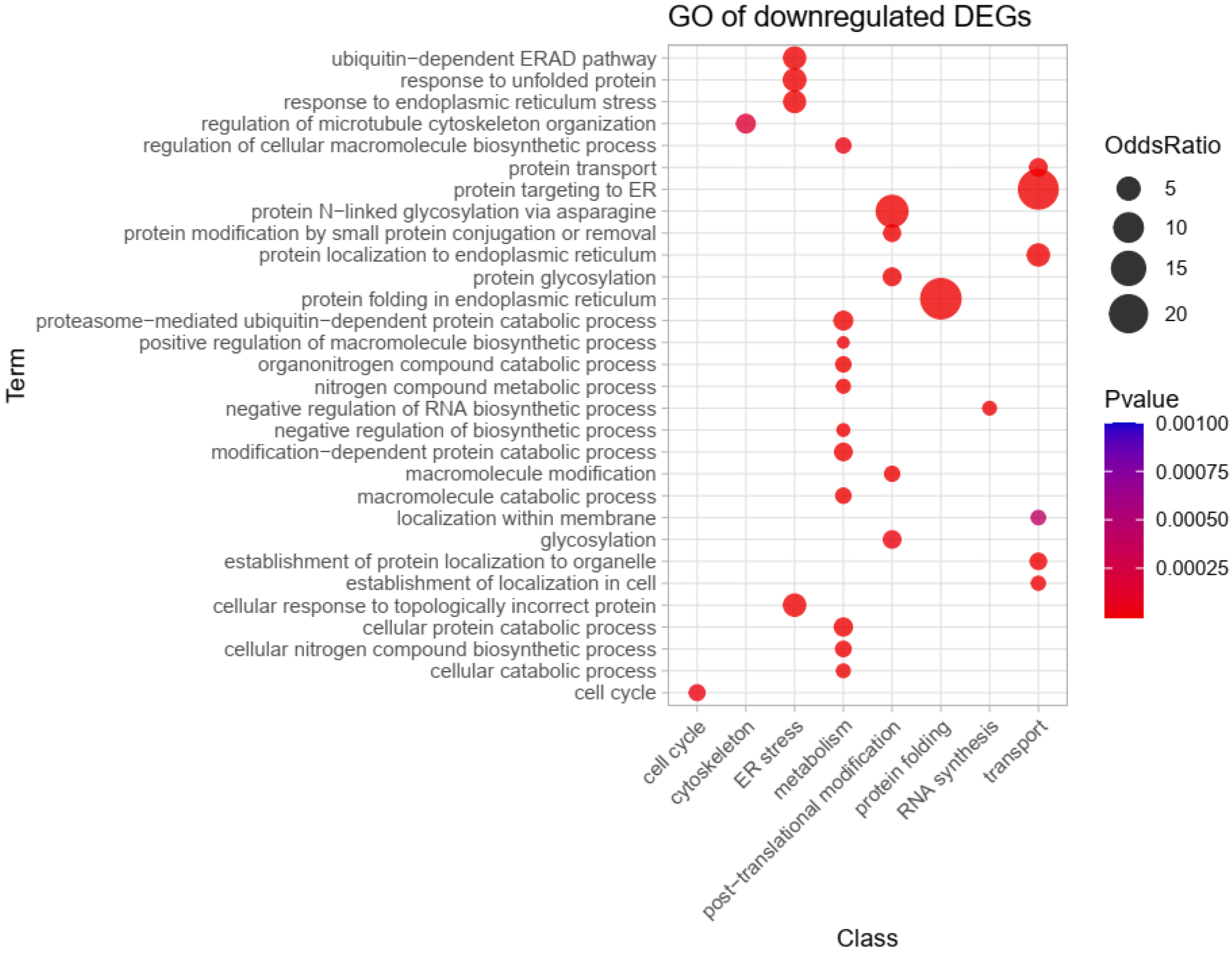
GO overrepresentation analysis of downregulated pathways.

**Fig EV3.**
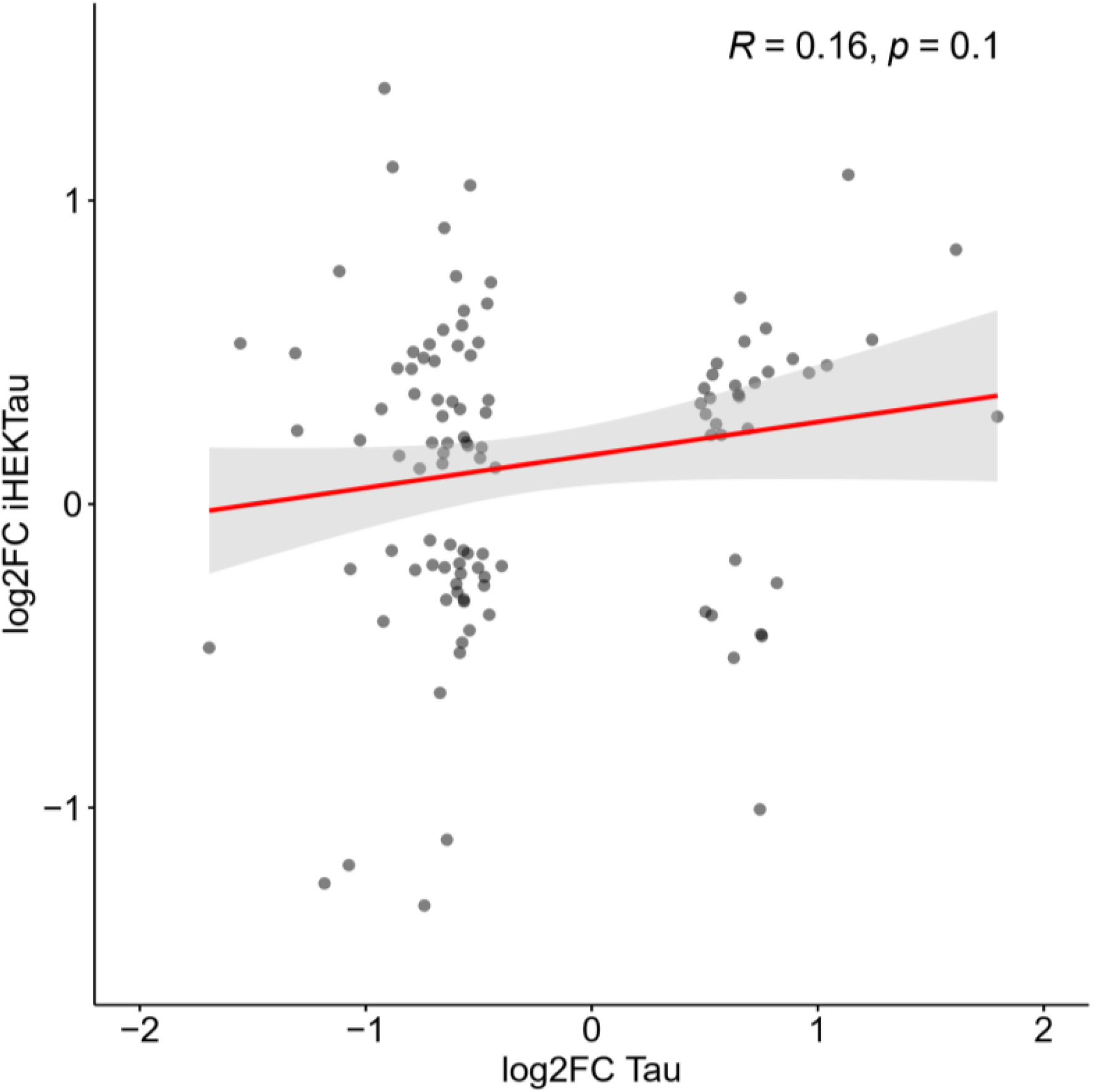
Scatter plot representing the log2FCs of genes in proAD-mimic cells (x-axis) and iHEK cells expressing Tau from dataset PRJNA744518 (Montalbano et al., 2021).

**Table EV1.** GO pathways identified in Darkgrey module are reported

**Table EV2.** GO pathways identified in Pink module are reported

